# Machine Learning Reveals Genetic Modifiers of the Immune Microenvironment of Cancer

**DOI:** 10.1101/2022.12.13.520300

**Authors:** Bridget Riley-Gillis, Shirng-Wern Tsaih, Emily King, Sabrina Wollenhaupt, Jonas Reeb, Amy R. Peck, Kelsey Wackman, Angela Lemke, Hallgeir Rui, Zoltan Dezso, Michael J. Flister

## Abstract

Heritability in the immune tumor microenvironment (iTME) has been widely observed, yet remains largely uncharacterized and systematic approaches to discover germline genetic modifiers of the iTME still being established. Here, we developed the first machine learning approach to map iTME modifiers within loci from genome-wide association studies (GWAS) for breast cancer (BrCa) incidence and outcome. A random forest model was trained on a positive set of immune-oncology (I-O) targets using BrCa and immune phenotypes from genetic perturbation studies, comparative genomics, Mendelian genetics, and colocalization with autoimmunity and inflammatory disease risk loci. Compared with random negative sets, an I-O target probability score was assigned to the 1,362 candidate genes in linkage disequilibrium with 155 BrCa GWAS loci. Pathway analysis of the most probable I-O targets revealed significant enrichment in drivers of BrCa and immune biology, including the *LSP1* locus associated with BrCa incidence and outcome. Quantitative cell type-specific immunofluorescent imaging of 1,109 BrCa patient biopsies revealed that LSP1 expression is restricted to tumor infiltrating leukocytes and correlated with BrCa patient outcome (HR = 1.73, p < 0.001). The human BrCa patient-based genomic and proteomic evidence, combined with phenotypic evidence that *LSP1* is a negative regulator of leukocyte trafficking, prioritized *LSP1* as a novel I-O target. Finally, a novel comparative mapping strategy using mouse genetic linkage revealed *TLR1* as a plausible therapeutic candidate with strong genomic and phenotypic evidence. Collectively, these data demonstrate a robust and flexible analytical framework for functionally fine-mapping GWAS risk loci to identify the most translatable therapeutic targets for the associated disease.

## INTRODUCTION

The heritability of breast cancer (BrCa) is estimated to be ~30% (Peto and Mack 2000; Hartman et al. 2007; Rosman et al. 2007; Pharoah et al. 2002) and germline genetic modifiers have been implicated in most aspects of the disease, including incidence (Michailidou et al. 2013; Flister and Bergom 2018), age-of-onset (Rafiq et al. 2013), distant metastasis (Park et al. 2005), and survival (Shu et al. 2012; Morra et al. 2021; Escala-Garcia et al. 2019). Genome-wide association studies (GWAS) have identified >170 genetic loci to date (Zhang et al. 2020) and hundreds of genes have been prioritized as candidates by genetic fine-mapping strategies, including functionally validated modifiers of tumor intrinsic phenotypes (Fachal et al. 2020). In contrast, germline genetic modifiers of BrCa that function through the tumor microenvironment (TME) remain poorly understood (Flister and Bergom 2018).

The immune system is a key component of the TME, and several recent studies have highlighted germline genetic contribution to antitumor immunity (Flister and Bergom 2018; Sayaman et al. 2021; Shahamatdar et al. 2020; Pagadala et al. 2021; Thorsson et al. 2018; Lim et al. 2018). A GWAS of the Cancer Genome Atlas (TCGA) pan-cancer cohort revealed heritability (~15-20%) across 33 traits related to tumor infiltrating leukocytes, including key modulators of antitumor immunity (e.g., IFN, STING, and cytotoxic T-lymphocytes) (Sayaman et al. 2021). Likewise, a separate TCGA pan-cancer analysis found multiple gene-level associations with 16 immune-related traits, including overlapping GWAS candidates for autoimmune diseases (Shahamatdar et al. 2020). These data, combined with evidence that similar immune traits predict response to anticancer immunotherapies and overall patient survival (Pagadala et al. 2021; Thorsson et al. 2018; Lim et al. 2018), suggest that germline genetic modifiers of the immune TME (iTME) likely impact cancer incidence, progression, and outcome.

Despite the evidence that heritable variation in antitumor immunity impacts BrCa (Flister and Bergom 2018), attempts to interrogate germline genetic modifiers of the iTME remain challenging for several reasons. Firstly, well-powered BrCa GWAS (e.g., ~118,000 cases in BCAC) do not collect iTME phenotypes (Zhang et al. 2020; Fachal et al. 2020), whereas smaller GWAS (e.g., ~9,000 cases in TCGA) with iTME phenotypes are underpowered for robust genome-wide genetic association (Sayaman et al. 2021; Shahamatdar et al. 2020; Pagadala et al. 2021; Thorsson et al. 2018; Lim et al. 2018). Secondly, genome-wide genetic perturbation datasets for functionally annotating candidates have only recently become available (Boehm et al. 2021; Stark et al. 2006) and functional genomics data have yet to be systematically incorporated into fine-mapping strategies for GWAS loci. Finally, in some cases, characterization of iTME modifiers will require comparative analysis using models with a fully intact iTME and yet there is a dearth of genetically diverse syngeneic cancer models for testing iTME modifiers.

The objective of this study was to develop an analytical framework for using human BrCa GWAS as a roadmap for translating iTME modifiers with the greatest therapeutic potential for antitumor immunity. To achieve this objective, a random forest model was trained on a positive set of immune-oncology (I-O) targets using BrCa and immune phenotypes from genetic perturbation studies, comparative genomics, Mendelian genetics, and colocalization with autoimmunity and inflammatory disease risk loci. When applied to 1,362 candidate genes in linkage disequilibrium (LD, *r^2^* > 0.6) with 155 human BrCa GWAS loci, several novel I-O targets were identified, including *LSP1* and *TLR1*. These data, combined with genomic, transcriptomic, proteomic, and phenotypic evidence, collectively demonstrate that fine-mapping with systematic integration of functional data (i.e., functional fine-mapping) is a powerful approach for translating GWAS risk loci to novel therapeutic targets.

## RESULTS

### Random forest modeling of I-O target probability for breast cancer GWAS candidates

Recent GWAS in TCGA have demonstrated that germline genetic modifiers of the iTME are prevalent in human cancer, including multiple candidates that are current I-O therapeutics (Sayaman et al. 2021; Shahamatdar et al. 2020; Pagadala et al. 2021; Thorsson et al. 2018; Lim et al. 2018). Although these GWAS are relatively small and underpowered for candidate gene discovery, the observations suggest that larger cancer GWAS might provide a roadmap for I-O targets with increased translational potential due to the existing association with human malignancy. To this end, colocalization analysis of loci was performed using multiple large publicly available GWAS (N=809,871 unique individuals) for BrCa (n=155 loci with 452 lead SNPs curated in this study) and autoimmunity (n=4,076 loci), which confirmed that heritable immunity is a prevalent component of BrCa risk (48% of BrCa loci, *p*<0.0001) (**Figure 1A** and **Table S1-S3**). However, due to the paucity of immunophenotypic data from these BrCa GWAS, identifying iTME modifiers is not possible without integrating immune phenotypes from other sources, such as Mendelian genetics, comparative genomics, and genetic perturbation studies. The objective of this study was to develop a machine learning approach that would identify the most probable antitumor immune modifiers from BrCa GWAS loci by learning the shared properties of known IO targets (i.e., the positive set) compared to a negative set of proteins.

**Figure 1.**
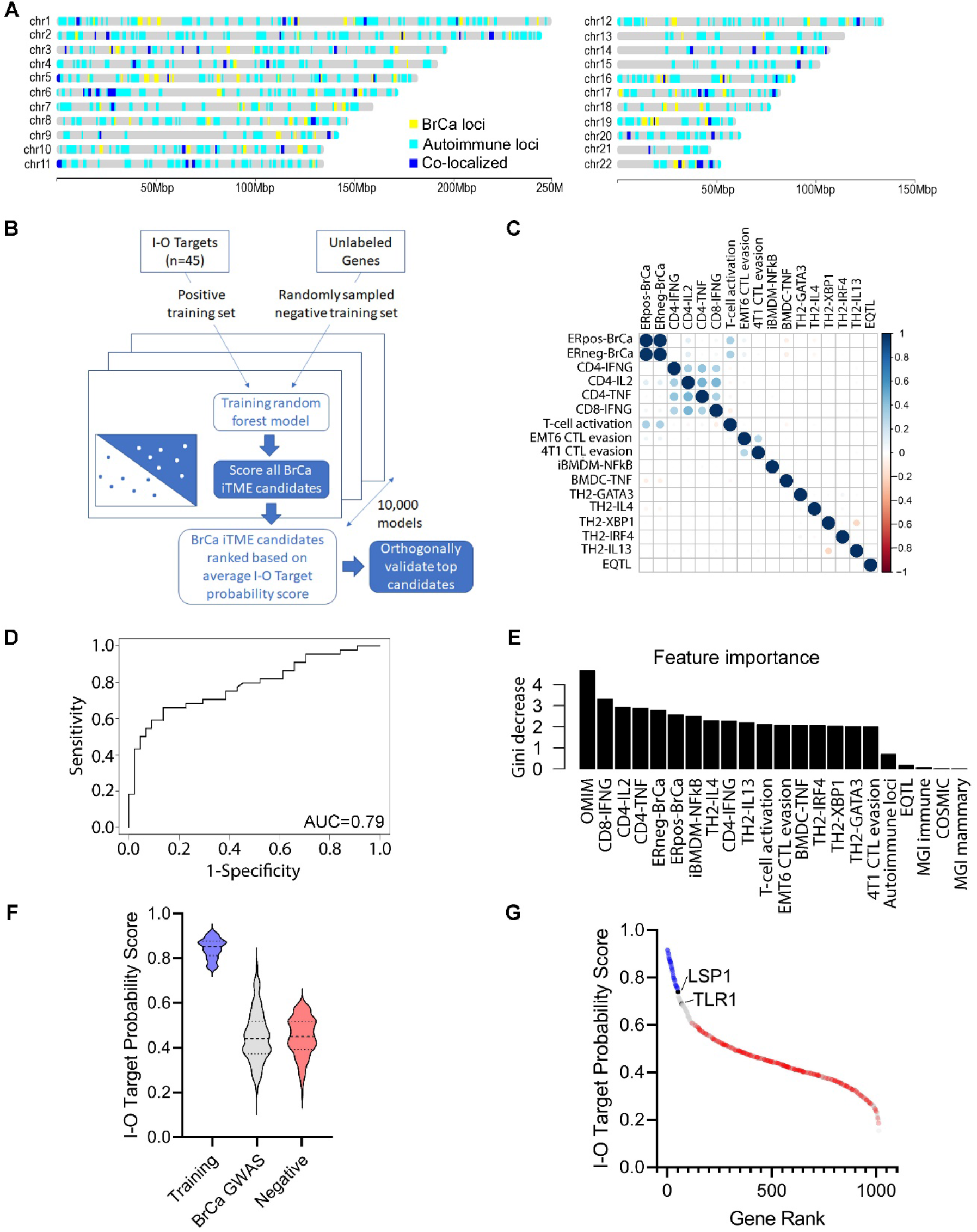
Random forest modeling of I-O target probability for breast cancer GWAS candidates. (**A**) global alignment of GWAS risk loci for BrCa (n = 155 loci with 452 lead SNPs curated in this study) and autoimmunity (n = 4,076 loci from Open Targets Genetics) revealed significant colocalization (48% of BrCa loci, *p*<0.0001). (**B**) Overview of the I-O target probability predictions using a positive training set of genes with preclinical and clinical evidence as I-O therapies. A size-matched negative set was generated from random sampling without replacement of all other genes and after excluding the positive set. Random forest models were then built (n = 10,000 models) using each of the random negative sets to generate predictions, followed by averaging across the 10,000 models to assign a I-O target probability score to 1,362 candidate genes in linkage disequilibrium (LD, *r^2^* > 0.6) with 155 human BrCa GWAS loci. (**C**) A correlation matrix of the quantitative variables revealed limited overlap between features. (**D**) Receiver operating characteristic (ROC) from leave-one-out cross validation using an independent negative set curated from the non-essential gene list and filtered on genes without reported expression in immune or breast cells. The prediction performance was reported as AUC (area under the curve) of 0.79. (**E**) The distribution of I-O target probability scores in the training set, BrCa candidates, and independent negative set. (**F-G**) Ranking of I-O target probability scores across the training set (blue), BrCa candidates (grey), and the independent negative set (red). Note the high probability rankings of the BrCa candidates, *LSP1* and *TLR1*.

We hypothesized that a random forest model that implemented *“easy ensemble” ensemble* (He and Garcia 2009) and *bagging* (Breiman 1996) would be well-suited to address the challenge of a limited set of true positives (n=45 curated I-O targets) and a larger set of unknown potential targets (i.e., all other genes). The resulting unbalanced positive-only learning task can be addressed by using negative training sets of equivalent size to the positive set and randomly sampling unique sets that exclude the positive set (Dezső and Ceccarelli 2020) (**Figure 1B**). Random forest models were then built using the averaged predictions over the 10,000 models to assign an I-O target probability score to the 1,362 BrCa candidates in LD (*r^2^* > 0.6) with 155 human BrCa GWAS loci. To capture unique aspects of iTME biology, the random forest model integrated features from complex genetics (**Table S1-S3**), Mendelian genetics (**Table S4**), genetic perturbation studies (**Table S5-S19**), comparative genomics (**Table S20-S21**), cancer drivers (**Table S22**), and eQTLs (**Table S23**). A Pearson’s correlation revealed that there was low covariance between the quantitative variables that were included as features in the random forest model (**Figure 1C**), suggesting that feature contributions to the model were largely independent. To assess model performance, a leave-one-out cross validation was performed using the positive set matched with a randomly sampled negative set of the same size, followed by receiver operator characteristic (ROC) area under the curve (AUC) analysis averaged over 100 models (**Figure 1D**). The mean decrease in the Gini metric was used to assess the relative importance of the features in the models, which revealed that Mendelian disease feature was the most predictive and followed closely by genetic perturbations and other features (**Figure 1E**). Combined these features performed expectedly well in predicting the training set (median score = 0.83) compared with the negative training (median score = 0.43) and the BrCa candidates (median score = 0.44) (**Figure 1F**). Among the BrCa candidates was a small group of outliers with I-O target probability scores approaching the training set, which included the immune modulators, *LSP1* and *TLR1* (**Figure 1G, Table S24**).

### *LSP1* is a novel I-O target associated with BrCa risk and outcome

*LSP1* is a negative regulator of leukocyte trafficking and activation (Jongstra-Bilen et al. 2000, 1), which has been replicated across multiple GWAS for for BrCa incidence (Michailidou et al. 2013, 2017; Easton et al. 2007; Michailidou et al. 2015; Shu et al. 2020) and outcome (Barrdahl et al. 2015; [CSL STYLE ERROR: reference with no printed form.]), as well as inflammatory and autoimmune diseases (Anderson et al. 2011; Liu et al. 2015; Chen et al. 2020; Kachuri et al. 2021; Astle et al. 2016), yet the therapeutic potential of *LSP1* is unknown. Based on the random forest analysis, the probability of *LSP1* as an I-O therapeutic ranked in the top 1% of BrCa GWAS candidates (8^th^ out of 1,362 candidates) (**Table S24**), suggesting that *LSP1* is an iTME modifier that shares the characteristics of successful I-O targets. Further, seven LD blocks (*r*^2^ > 0.6) localized to *LSP1* have been associated with BrCa incidence or outcome, including 5 LD blocks that correlated with altered LSP1 expression in leukocytes and 29 out of 78 SNPs (37%) within the LD blocks that were predicted to alter a functional motif (**Table S25**). Notably, 19 of the 29 functional SNPs (fSNPs) (66%) were predicted to disrupt canonical binding sites for transcriptional regulators of immune and inflammatory signaling (**Table S25**). For example, the most significantly BrCa-associated LD block (led by rs1973765) (Michailidou et al. 2017) contained three fSNPs that were predicted to disrupt *LSP1* regulatory regions, including binding sites for immune regulatory transcription factors, ASCL2 (rs4980389) and PLAG1 (rs588321) (**Figure 2A**), and eight SNPs correlated with pan-leukocyte expression of LSP1 (**Figure 2B**). Finally, *LSP1* expression in treatment-naïve tumor specimens from BrCa patients was largely restricted to the infiltrating leukocytes (Qian et al. 2020; Bassez et al. 2021) (**Figure 2C, 2D**) and high leukocyte-specific expression of *LSP1* correlated with shorter progression-free survival in univariate analysis of BrCa (HR = 1.73, 95% CI 1.29–2.33; p < 0.001) (**Figure 2E–2K**). Combined, these data establish that *LSP1* is a heritable iTME modifier that correlates with BrCa patient survival and closely resembles the characteristics of known I-O targets, which collectively suggest that *LSP1* is a potential I-O therapeutic target with high translational potential.

**Figure 2.**
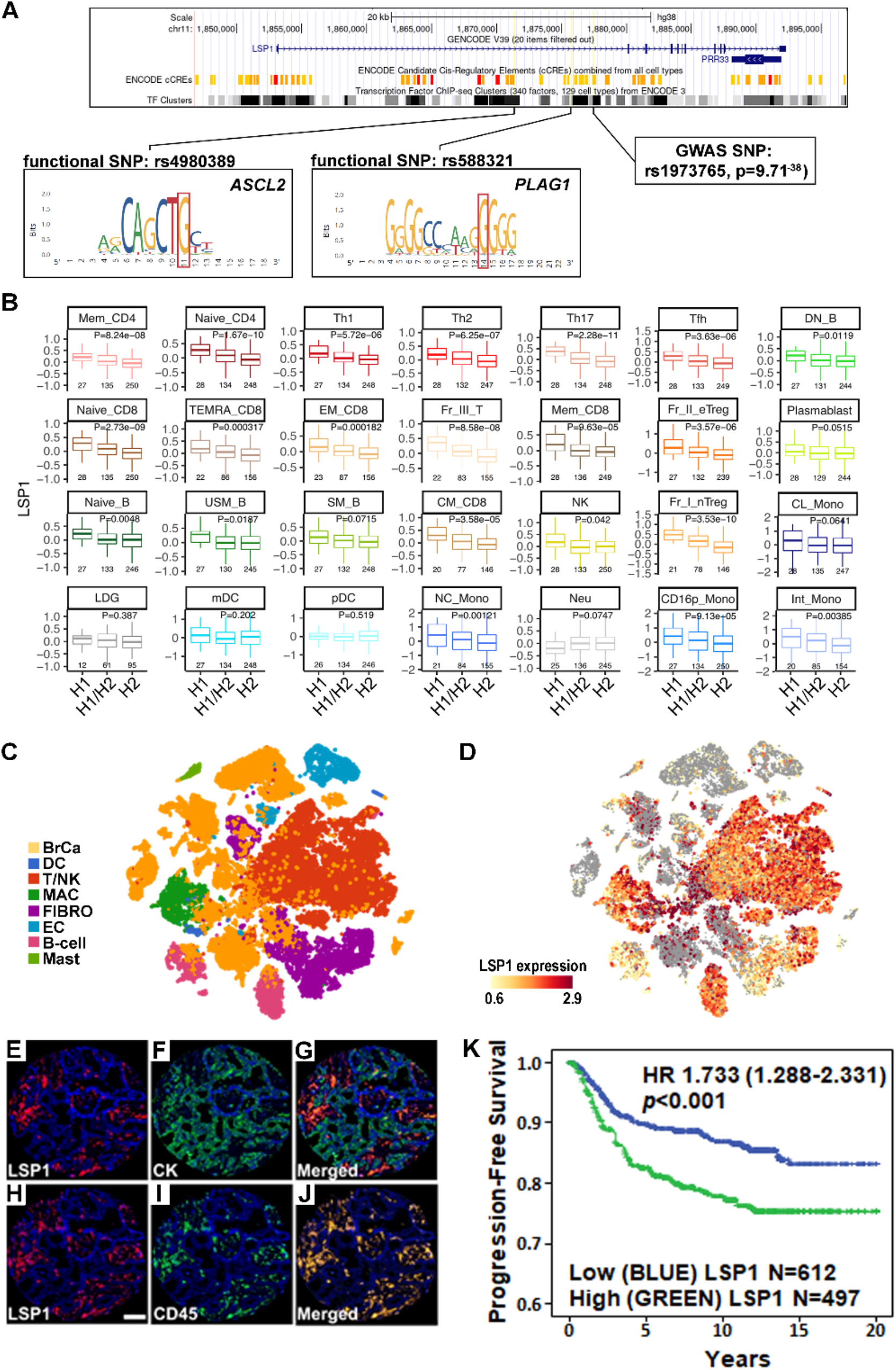
*LSP1* is a germline genetic iTME modifier of BrCa incidence and outcome. (**A**) Annotation of a BrCa-associated *LSP1* locus (rs1973765) in LD (*r^2^* < 0.6) with multiple fSNPs (e.g., rs4980389 and rs588321) and 17 eSNPs associated with pan-leukocyte expression of LSP1. (**C**) scRNAseq analysis of treatment naïve BrCa patient biopsies profiled for different cell populations BrCa cells (BrCa), dendritic cells (DC), T-cells and NK cells (T/NK), macrophages (MAC), fibroblasts (FIBRO), endothelial cells (EC), B-cells, and mast cells (MAST). (**D**) Expression of LSP1 detected by scRNAseq across the same treatment naïve BrCa patient biopsies. (**E-J**) Quantitative immunofluorescent imaging of LSP1 protein expression colocalized to tumor cells (pan-CK^+^) or infiltrating leukocytes (CD45^+^) in tumors of untreated BrCa patients. (**K**) Progression-free survival analysis based on CD45^+^ leukocyte-specific expression of LSP1 across tumors from 1,109 BrCa patients.

### A novel comparative mapping strategy in the iTME identifies *TLR1* as an I-O target

We recently demonstrated that genetic modifiers in the TME can be mapped using cancer xenograft host strains with distinct genetic backgrounds that differ cancer risk profiles (Flister et al. 2014, 2017). Because the xenograft host strain backgrounds vary, whereas the malignant tumor cells do not, any observed changes in tumor progression are due to genetic differences in the nonmalignant TME (Flister et al. 2014, 2017). However, a challenge in using human xenografts for TME mapping studies is that the iTME compartment is inaccessible due to the use of immunocompromised xenograft strains. Thus, we developed the first strategy for mapping iTME modifiers using syngeneic mouse models with different susceptibility to mammary cancer.

PWD/PhJ is an inbred wild-derived *Mus. m. musculus* strain that is genetically diverse from *Mus. m. domesticus* and therefore well suited for genetic trait mapping with common laboratory strains, such as C57BL/6J (Gregorová and Forejt 2000). To assess whether the genetic modifiers observed in the PWD/PhJ genome function through the host TME, the syngeneic C57BL/6J-derived E0771.LMB mammary tumor cell line was implanted into C57BL/6J (B6) homozygous female mice (n = 22) and PWD/PhJ x C57BL/6J (PWD.B6) F1 heterozygous female mice (n = 10). Because both host models contain at least one full copy of the C57BL/6J genome, the E0771.LMB cells are syngeneic with the B6 and PWD.B6 host strains (**Figure 3A**). At 17 days post-implantation, the E0771.LMB tumor burden in PWD.B6 mice (619 ± 147 mm^3^ and 0.5 ± 0.1g) was significantly less than B6 tumors (1,830 ± 188 mm^3^, *P* < 0.001 and 1.5 ± 0.1g, *P* < 0.001) (**Figure 3B, 3C**). Bulk RNAseq analysis of E0771.LMB tumors grown in B6 and PWD.B6 mice (n=4 per group) revealed 750 differentially expressed genes (DEGs) (**Table S26**) and gene set enrichment (GSE) suggested increased inflammation (**Figure 3D**) and leukocyte infiltration (**Figure 3E**) in PWD.B6 mice. Likewise, known antitumor immunomodulatory pathways were predicted to be highly activated in E0771.LMB tumors implanted in PWD.B6 mice compared with B6: IFNγ (pathway z-score = 6.7; p = 1.74^-24^), TNFα (pathway z-score = 6.2; p = 8.52^-21^), and IL1B (pathway z-score = 4.9; p = 1.29^-22^). To orthogonally test whether the PWD/PhJ and C57BL/6J genomic backgrounds differ in susceptibility to spontaneous mammary tumors driven by the MMTV-PyMT transgene, the latency of mammary tumors in homozygous B6-Tg(MMTV-PyMT) mice were compared with heterozygous PWD.B6-Tg(MMTV-PyMT) Fi female mice up to 20 weeks-of-age (**Figure 4A**). Compared with B6-Tg(MMTV-PyMT), the median tumor latency of PWD.B6-Tg(MMTV-PyMT) F_1_ female mice was significantly delayed by 5 weeks (p<0.001) (**Figure 4B**). No differences in MMTV-PyMT transgene expression in the mammary tumors of B6-Tg(MMTV-PyMT) and PWD.B6-Tg(MMTV-PyMT) mice was detected (**Figure 4C**), indicating that strain-dependent PyMT oncogene expression does not contribute to differences in the mammary tumor risk between B6-Tg(MMTV-PyMT) and PWD.B6-Tg(MMTV-PyMT) mice. Combined with the reduced growth potential of E0771.LMB mammary tumors in PWD.B6 mice, these data suggest that one or more modifiers impact the host iTME to suppress mammary cancer.

**Figure 3.**
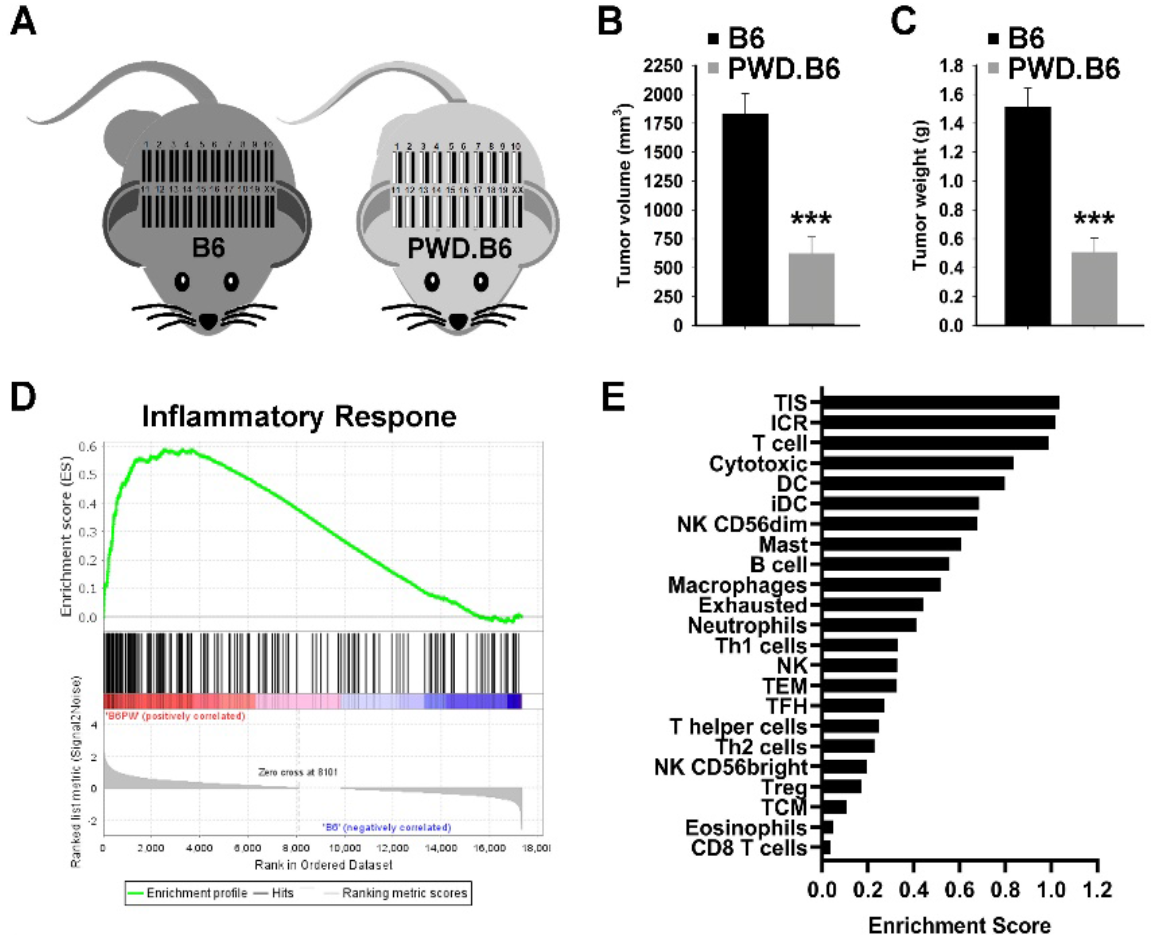
Comparative mapping of iTME modifiers in mouse and human BrCa. (**A**) Schematic representation of syngeneic tumor transplant experiments in homozygous C57BL/6J (B6) mice and PWD/PhJ x C57BL/6J (PWD.B6) F_1_ heterozygous mice. E0771.LMB tumor volumes (**B**) and weights (**C**) at 17 days postimplantation in B6 homozygous female mice (n = 22) and PWD.B6 F_1_ heterozygous female mice (n = 10). Enrichment of differentially expressed genes in inflammatory pathways (**D**) and immune cell infiltrates (**E**) assessed by GSEA of bulk transcriptomic data from E0771.LMB tumors grown in a B6 and PWD.B6 Fi mice (n = 4 per group).

**Figure 4.**
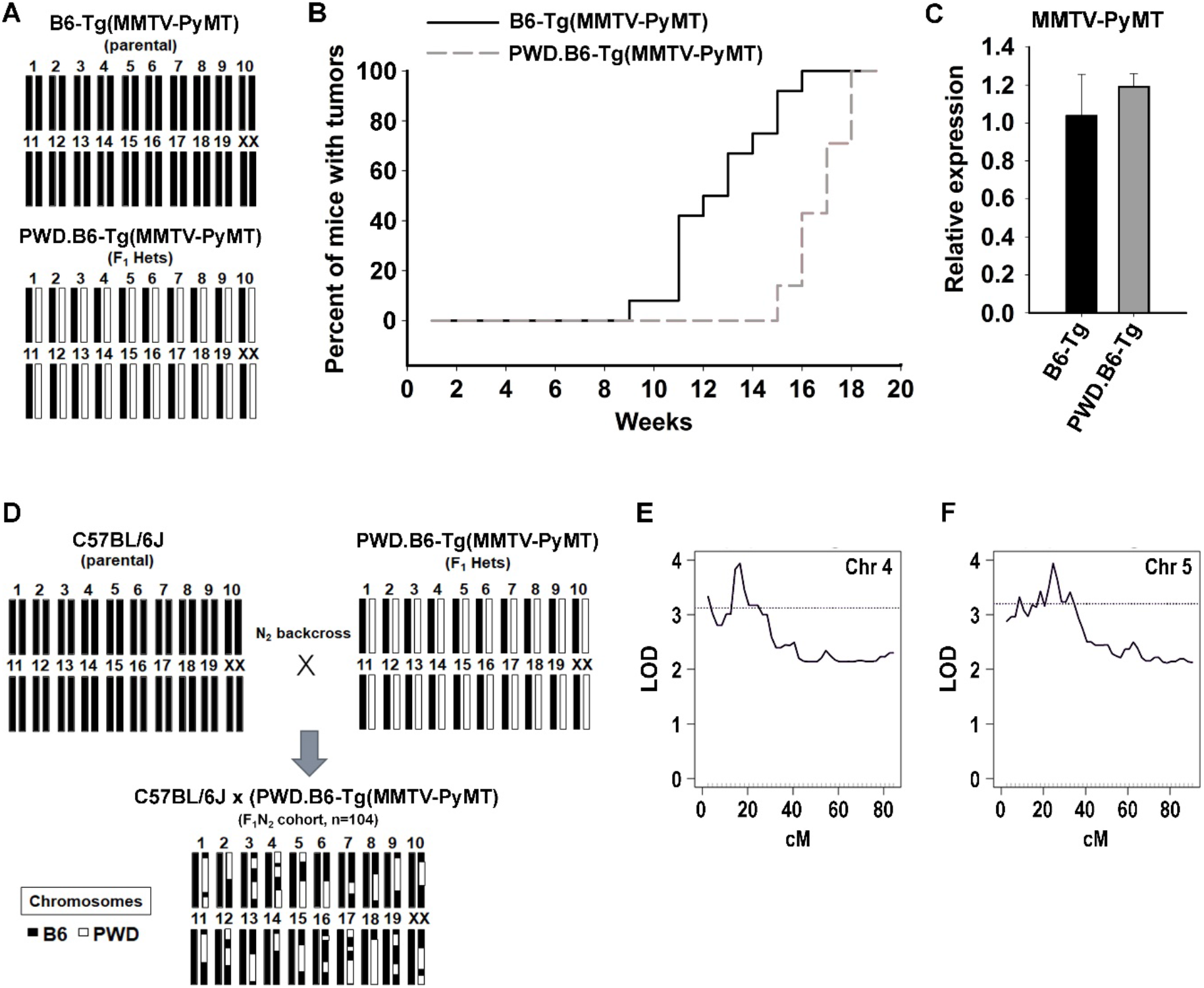
Comparative mapping of iTME loci in the PWD/PhJ mouse background. (**A**) Schematic representation of the genomic backgrounds of homozygous B6-Tg(MMTV-PyMT) mice (black bars) and heterozygous PWD.B6-Tg(MMTV-PyMT) F_1_ mice (white bars). (**B**) Incidence and latency of mammary tumorigenesis in homozygous B6-Tg(MMTV-PyMT) mice and heterozygous PWD.B6-Tg(MMTV-PyMT) F_1_ mice. (**C**) Relative expression of MMTV-PyMT in tumors extracted from homozygous B6-Tg(MMTV-PyMT) mice and heterozygous PWD.B6-Tg(MMTV-PyMT) F_1_ mice (n = 4 per group). (**D**) Schematic representation of the C57BL/6J x (PWD.B6-Tg(MMTV-PyMT)) N2 backcross (n = 104). (**F**) Linkage of chromosome 4 with mammary tumor latency when jointly considering the chromosome 5 QTL. (**G**) Linkage of chromosome 5 with mammary tumor latency when jointly considering the chromosome 4 QTL.

Interval mapping of a C57BL/6J x (PWD.B6-Tg(MMTV-PyMT)) N2 backcross (n = 104) was used to localize genetic modifier(s) of mammary tumorigenesis (**Figure 4D**). A genome-wide single locus scan revealed suggestive PWD/PhJ loci that increased mammary tumor latency at chr4 (peak: 34.03 Mb; support interval: 4.68 - 55.84 Mb) and chr5 (peak: 44.93 Mb; support interval: 15.97 - 66.36 Mb). When jointly considered, the PWD/PhJ alleles within the chr4 and chr5 loci were significantly associated with increased tumor latency (LOD = 3.9; p = 1 x 10^-4^) (**Figure 4E, 4F**). The mouse chr4 locus was not syntenic to any reported human BrCa GWAS locus, whereas the chr5 locus was syntenic to one human GWAS locus spanning three candidates (*TLR1, WDR19*, and *TMEM156*) that were associated with BrCa risk (Zhang et al. 2020; Fachal et al. 2020). Of the candidates, the *Tlr1* allele had the strongest genotypic and phenotypic evidence, including elevated expression and pathway activation in E0771.LMB tumors implanted in the PWD.B6 genetic background (**Figure 5A** and **Table S26**), as well as multiple potentially damaging coding variants in the PWD/PhJ background (**Tables S27-S28**). Additionally, *TLR1* had the highest I-O target probability score (0.69) of the candidates, colocalized with autoimmune loci (Kichaev et al. 2019), and was functionally linked with immune phenotypes ([CSL STYLE ERROR: reference with no printed form.]; Shifrut et al. 2018; Lawson et al. 2020; [CSL STYLE ERROR: reference with no printed form.]; Covarrubias et al. 2020) (**Figure 5B, 5C**). In contrast, no intrinsic BrCa phenotypes were observed for *TLR1* (**Figure 5B, 5C**). The *TLR1* locus that colocalized with risk loci for BrCa (rs6815814, p=6.13^-13^) (Michailidou et al. 2017) and autoimmunity (rs5743618, p=4^-114^) (Kichaev et al. 2019) was also in LD (r^2^ > 0.6) with multiple fSNPs predicted to disrupt transcriptional regulatory regions (rs5743565 and rs11722889) (**Figure 5D**) and multiple eSNPs that correlated with expression across various human leukocyte cell types (**Figure 5E-H**). Collectively, these data suggest that *TLR1* is a novel iTME modifier of BrCa and a molecular target with high translational potential due to the existing association with human BrCa.

**Figure 5.**
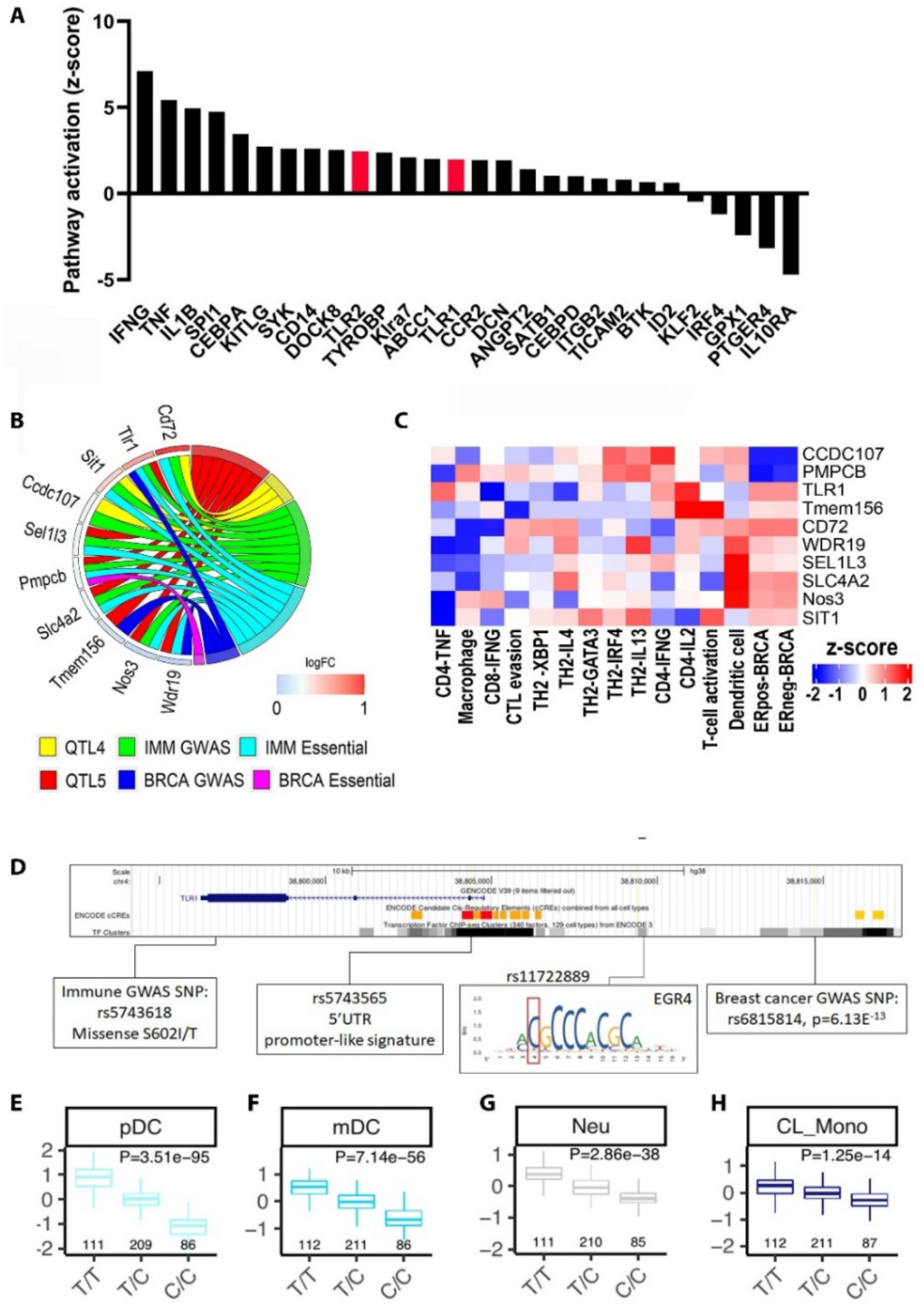
Evidence of *TLR1* as an iTME modifier of BrCa. (**A**) Pathway activation (z-score) predicted from bulk transcriptomic analysis of E0771.LMB tumors grown in a B6 and PWD.B6 F_1_ mice (n = 4 per group). Significant activation of the top iTME candidate pathway, Tlr1/2, is highlighted in red. (**B**) Distribution of intrinsic BrCa phenotypes and extrinsic iTME phenotypes across the candidates overlapping with human GWAS loci. (**C**) Heatmap of functional fine-mapping scores using CRISPR screening data related to intrinsic BrCa phenotypes and extrinsic iTME phenotypes. (**D**) Example of a pleiotropic *TLR1* haplotype that colocalizes with risk loci for BrCa (rs6815814) and autoimmunity (rs5743618), as well as multiple fSNPs (rs5743565 and rs11722889) within transcriptional regulatory regions of the *TLR1* locus. Examples of eSNP (rs5743565) for TLR1 across multiple human leukocyte cell types, including plasmacytoid dendritic cells (pDC) (**E**), Myeloid dendritic cells (mDC) (**F**), neutrophils (Neu) (**G**), and classical monocytes (**H**).

## DISCUSSION

Germline genetics contribute to variability in the tumor-infiltrating immune system (referred to as iTME) (Flister and Bergom 2018; Sayaman et al. 2021; Shahamatdar et al. 2020; Pagadala et al. 2021; Thorsson et al. 2018; Lim et al. 2018), which is clinically relevant to disease progression and response to anticancer immunotherapies (Pagadala et al. 2021; Thorsson et al. 2018; Lim et al. 2018). However, most germline genetic modifiers of the iTME remain uncharacterized due to the lack of well-powered GWAS that capture iTME phenotypes. Further compounding the challenges with mapping iTME modifiers is the limited integration of functional and comparative data for iTME-relevant phenotypes into current fine-mapping strategies for GWAS. To overcome these challenges, we developed a novel machine learning approach that used genomic, transcriptomic, proteomic, and phenotypic evidence to identify the BrCa GWAS candidate genes that most closely resemble features of a positive set of I-O therapeutic targets. This novel approach prioritized multiple BrCa candidate genes that resembled novel I-O therapeutics, such as *LSP1* and *TLR1*, which were orthogonally validated by human BrCa disease association and comparative modeling, respectively. Collectively, these data demonstrate for the first time that germline genetic iTME modifiers within BrCa GWAS loci are pervasive and fine-mapping these loci with systematic integration of functional data is a powerful approach for translating GWAS risk loci to novel therapeutic targets.

### Approaches to mapping heritability in the iTME

Although the heritability of germline genetic modifiers of the iTME has long been suspected (Flister and Bergom 2018), it has only recently been measured using TCGA cohorts that uniquely have both patient genotypes and immune phenotypes inferred from bulk RNAseq (Sayaman et al. 2021; Shahamatdar et al. 2020; Pagadala et al. 2021; Thorsson et al. 2018; Lim et al. 2018). The estimated heritability of iTME modifiers from these studies approached 20% of variance for some traits (Sayaman et al. 2021; Pagadala et al. 2021), excluding highly polymorphic regions with much higher estimates of heritability, such as the HLA and KIR regions (Pagadala et al. 2021). The collective estimates of iTME heritability from TCGA (Sayaman et al. 2021; Pagadala et al. 2021) match the estimated heritability of immune traits in healthy individuals by larger GWAS (Astle et al. 2016; Orrù et al. 2013) and twin studies (Brodin et al. 2015), suggesting that the heritability of immunity is consistent in normal and malignant settings. However, despite comparable levels of heritable immunity in normal (Orrù et al. 2013) and malignant settings (Sayaman et al. 2021; Pagadala et al. 2021), only a handful of iTME candidates were detected by GWAS in TCGA pan-cancer cohort (~9,000 patients) as compared with >2,700 modifier loci of immune traits in a much larger cohort of >500,000 healthy individuals (Astle et al. 2016). Together, these observations indicate that GWAS in TCGA is currently underpowered to detect all but the most penetrant iTME modifier loci, and larger cohorts with matched germline and RNAseq data are needed to replicate and expand the catalog of germline iTME modifiers (Shahamatdar et al. 2020). The inadequate power to detect most iTME modifiers by GWAS in TCGA is further compounded by the limitations of deconvoluting cell type-specific signals within bulk transcriptomics data (Newman et al. 2015). Thus, until well-powered cancer patient datasets expand with corresponding immune phenotypes, identifying iTME modifiers is not possible without integrating immune phenotypes from other sources, such as Mendelian genetics, comparative genomics, and genetic perturbation studies.

To our knowledge, this study was the first to develop a machine learning approach to map the most probable antitumor immune modifiers from BrCa GWAS loci by learning the shared properties of known I-O targets (i.e., the positive set) without a labeled set of negative controls. In this setting, the learner only has access to a small set of positive examples and a large set of unlabeled data, because outside the known I-O targets there are many genes with unknown potential as I-O therapeutic targets. This type of machine learning problem is referred to as *Positive Unlabeled* (PU) (Cerulo et al. 2010; Elkan and Noto 2008) and it is compounded by the risk of overfitting when the sizes of the positive and unlabeled sets are unbalanced (He and Garcia 2009). To overcome this problem, we developed a machine learning approach that combines *easy ensemble* (He and Garcia 2009) and *bagging* (Breiman 1996) to balance training sets by randomly sampling negative sets that are size-matched to the positive set (Dezső and Ceccarelli 2020). Random forest models (n=10,000) were then generated using the positive set and each non-overlapping negative set from the unlabeled data to generate an averaged probability score across all models (Dezső and Ceccarelli 2020). By using known I-O targets as the positive training set, we hypothesized that this approach could be used to prioritize germline genetic modifiers of the iTME with the greatest probability as novel I-O therapeutic targets. Because this machine learning approach prioritizes candidates associated *a priori* with a disease of interest, the candidates that most closely resemble successful therapeutic targets are also likely to have high translational potential. Finally, this machine learning framework is flexible in that it requires only a positive set for training (i.e., known targets for any disease or phenotype) and can integrate genotypic, phenotypic, and functional data from independent sources to overcome the current limitations of disease association studies.

### BrCa iTME candidates with potential as I-O targets

*LSP1* and *TLR1* were highly ranked within the top 1-3% as two of the most probable I-O targets among BrCa candidate genes. *LSP1* was also distinct from other iTME candidates in that it has been associated with BrCa outcome (Barrdahl et al. 2015; [CSL STYLE ERROR: reference with no printed form.]) in addition to incidence of BrCa (Michailidou et al. 2013, 2017; Easton et al. 2007; Michailidou et al. 2015; Shu et al. 2020) and autoimmunity (Anderson et al. 2011; Liu et al. 2015; de Lange et al. 2017; Jostins et al. 2012), suggesting that *LSP1* is a novel germline iTME modifier of BrCa incidence and disease progression. Moreover, cell-type specific detection of LSP1 in BrCa tumor specimens revealed that LSP1 expression is predominantly restricted to infiltrating leukocytes and was significantly correlated with better outcome. Based on the observation that *Lsp1*-deficient mice exhibit increased leukocyte motility (Jongstra-Bilen et al. 2000) and enhanced response to immune checkpoint blockade (ICB) (Kwon et al. 2020), we hypothesize that *LSP1* is a negative regulator of antitumor immunity. This hypothesis is further supported by the putative role of *LSP1* in neutrophil actin dysfunction (NAD, OMIM 257150), a rare immunological disorder characterized by early onset recurrent infections (Boxer et al. 1974; Coates et al. 1991; Southwick et al. 1988). These data, combined with genetic evidence that the *LSP1* locus is associated with altered LSP1 expression in leukocytes and BrCa-specific outcomes, suggest that *LSP1* is a novel iTME modifier of BrCa incidence and disease progression. *TLR1* was also a notable example of a GWAS candidate for both BrCa (Fachal et al. 2020; Michailidou et al. 2017) and immune phenotypes (Kichaev et al. 2019; Sakaue et al. 2021; Jonsson et al. 2017; Mayerle et al. 2013; Johansson et al. 2019; Han et al. 2020; Ferreira et al. 2020), which were confirmed by comparative mapping and GEMM phenotyping (Alexopoulou et al. 2002). TLR1 is a key regulator of innate immune response to DAMPs released by cancer cells (Urban-Wojciuk et al. 2019) and TLR1/2 agonists have been shown to enhance efficacy of ICB via activation of CTLs (Sharma et al. 2019, 2; [CSL STYLE ERROR: reference with no printed form.]). Combined with the evidence that *LSP1* and *TLR1* both modulate ICB therapies in experimental models (Sharma et al. 2019, 2; [CSL STYLE ERROR: reference with no printed form.]; Kwon et al. 2020), our data suggesting that *LSP1* and *TLR1* are likely to be patient-relevant targets for modulating antitumor immunity.

### Challenges and future strategies for mapping heritability in the iTME

Despite the evidence of iTME heritability, analytical approaches to identify germline genetic modifiers of the iTME are limited and there are many unresolved questions. One challenge is that iTME candidates are likely to have complex interactions across multiple molecular pathways, cell types, and physiological functions (Flister and Bergom 2018). It is also highly plausible that some pleiotropic iTME modifiers impact both BrCa cells and infiltrating leukocytes. Further confounding the ability to characterize iTME candidates is the observation that multiple causative genes can co-segregate at the same GWAS locus (Flister et al. 2013). A single genetic locus could therefore elicit complex biological changes across multiple cell types and the combined effects are ultimately manifested at the phenotypic level. Likewise, seemingly unrelated iTME modifiers that are not connected at the molecular level might interact at the cellular or tissue levels by modifying the density or physiological efficacy of cellular mediators within the tumor. For example, the phenotypic effects of a genetic modifier of infiltrating leukocyte function (e.g., *TLR1*) might be dampened or amplified in a patient that has coinherited a modifier of leukocyte trafficking (e.g., *LSP1*).

The challenges to disentangling the complexities of the host iTME will likely require further development of robust experimental and analytical tools for assessing germline genetic heritability in the iTME. Here, we took the initial step towards developing a machine learning approach identify the most likely iTME candidates within 1,362 genes in LD (*r^2^* > 0.6) with 155 human BrCa GWAs loci that resemble known I-O targets. To our knowledge, this was the first systematic analysis of BrCa GWAS loci to characterize iTME modifiers using the growing catalog of functional genomics datasets for both oncology and immunology. Intriguingly, this analysis revealed a prevalence of BrCa GWAS candidates (39%) with phenotypic evidence in leukocyte cell types, which also agrees with the significant overlap of 48% (*p*<0.0001) of BrCa GWAS risk loci with autoimmune disease loci. Notably, not all candidate phenotypes were observable in *ex vivo* phenotypic screens with isolated cell types, yet perturbation of some of these candidates in the *in vivo* and patient settings revealed strong phenotypic evidence in the iTME. For example, text mining patient data in OMIM was the most predictive feature in the random forest model, which could be attributed to disease phenotypes only observed in patients, such as the association of *LSP1* with the rare neutrophil actin dysfunction disorder (NAD, OMIM 257150). Experimental mapping using the first genetically diverse *in vivo* preclinical models for studying iTME heritability (C57BL/6J and PWD/PhJ), also enabled the independent replication of *TLR1* as a plausible germline genetic modifier of the iTME. Thus, future studies to expand the repertoire of genotypic, phenotypic, and functional data from preclinical and clinical datasets are likely to facilitate the discovery of additional germline genetic modifiers of the iTME that impact antitumor immunity and response to immunotherapies.

There also remains a need to develop approaches to directly measure the germline genetic contribution to the iTME in patients, which will further enable discovery of novel I-O targets with high translational potential. One possible approach would be to combine germline genotyping with single-cell RNA expression profiling (i.e., scRNAseq) of dissociated patient tumor biopsies to identify iTME eQTLs, as has been demonstrated elsewhere in circulating leukocytes isolated from healthy individuals (van der Wijst et al. 2018; Nathan et al. 2022). However, a major drawback of scRNAseq profiling using dissociated tissues is the inability to capture spatial orientation of profiled cells within the tissue. Spatial transcriptomics was recently developed to enable spatially resolved genome-wide gene expression at the near single-cell level (Andersson et al. 2021), yet both scRNAseq and spatial transcriptomics are cost prohibitive and not scalable to population level studies. Based on recent advances in tissue microarray technologies (Peck et al. 2016) and quantitative immunofluorescent imaging (Tran et al. 2010), we propose an alternative framework for iTME eQTL mapping at the protein level, using multiplex immunofluorescent assays to correlate cell-type specific protein expression with patient genotypes. The capacity of this novel approach could be expanded using high density tissue microarrays and quantitative immunofluorescent imaging, which offers highly sensitive and spatially resolved detection of protein expression at the cellular and subcellular levels.

### Conclusions

This study was the first to systematically map germline genetic modifiers of the iTME of human BrCa that most closely resembled the characteristics of known I-O targets, which enabled the discovery of *LSP1* and *TLR1* as novel iTME modifiers with therapeutic potential. This machine learning framework is flexible in that it required only a positive set for training and could integrate genotypic, phenotypic, and functional data from independent sources to overcome the current limitations of disease association studies. We orthogonally replicated the iTME-specific associations of *LSP1* and *TLR1* to BrCa risk using patient tumor biopsies and comparative mapping, respectively. Importantly, because this machine learning approach prioritizes candidates that were first identified by human BrCa risk association, we postulate that there is greater likelihood that *LSP1* and *TLR1* that will be translatable and clinically relevant to BrCa. Combined with our findings, these targets were also independently shown to modify responses to IBC in experimental cancer models (Sharma et al. 2019, 2; [CSL STYLE ERROR: reference with no printed form.]; Kwon et al. 2020). Thus, we conclude that *LSP1* and *TLR1* are likely to be patient-relevant targets for modulating antitumor immunity. Finally, these data demonstrate a robust and flexible analytical framework for functionally fine-mapping GWAS risk loci to identify the most translatable therapeutic targets for the associated disease.

## METHODS

### Human breast cancer GWAS candidates

Summary statistics were obtained from the Breast Cancer Association Consortium (BCAC) (Michailidou et al. 2017), UK Biobank (Bycroft et al. 2018), and FinnGen (https://www.finngen.fi/fi), a public-private partnership research project combining imputed genotype data generated from newly collected and legacy samples of Finnish biobanks and digital health record data from Finnish health registries (https://www.finngen.fi/en). The FinnGen data freeze 7 cohort included 309,154 individuals had been analyzed for over 3,000 endpoints, including incidence of BrCa and autoimmune diseases. The UKBB is a large and populationbased prospective cohort of approximately 500,000 participants aged 40-69 years recruited between 2006 and 2010 in the United Kingdom (Bycroft et al. 2018). The BCAC, FinnGen, and UKBB cohorts have a combined total of 152,059 BrCa cases and 657,812 controls of mostly European descent (**Table S1-S2**). Candidate gene mapping of the summary statistics was performed using FUMA (Watanabe et al. 2017), followed by positional mapping all SNPs in LD (*r*^2^ > 0.6) with the reported lead SNP and mapping genes to SNPs within 10Kb of the nearest gene using ANNOVAR. eQTL mapping were performed for all SNPs using the eQTL Catalogue (Kerimov et al. 2021), DICE (Schmiedel et al. 2018), GTEx (v8) (THE GTEX CONSORTIUM 2020), and eQTLGen (Võsa et al. 2021). LD blocks within select loci (e.g., *LSP1* and *TLR1*) were further characterized using Ensembl Variant Effect Predictor (VEP) (McLaren et al. 2016), ImmuNexUT (Ota et al. 2021), GWAS Catalog (MacArthur et al. 2017), and RegulomeDB (Boyle et al. 2012) to annotate SNPs overlapping function motifs (fSNPs).

### Colocalization of human breast cancer and autoimmunity risk loci

Autoimmunity risk variants were obtained from Open Targets Genetics version 6 (Mountjoy et al. 2021). Studies of autoimmune traits were identified using mapped Experimental Factor Ontology terms belonging to the class ‘autoimmune disease’ (Malone et al. 2010). Independent lead SNPs from BrCa loci were obtained as described below and are found in **Table S3**. Locus boundaries shown in the figure, plotted using the chromoMap R package (Anand and Rodriguez Lopez 2022), were constructed using 250 kb windows on either side of a lead SNP. To test for significant overlap between BrCa loci and autoimmune loci, each of the 155 BrCa loci was randomly assigned a uniformly distributed location on the genome with 10,000 permutations. The distance between the BrCa locus and the nearest autoimmune lead SNP was computed for each set of randomized BrCa locus, and this was used to compute the number of BrCa loci within 500 kb of an autoimmune lead SNP. Results are insensitive to the choice of summary statistic: the mean and median distance, the proportion BrCa loci containing a lead SNP for an autoimmune disease, the proportion of BrCa loci having an autoimmune SNP within varying distances from the locus up to 10 Mb similarly resulted in random distributions that were significantly different from observed values, in most cases without overlap.

### Random forest modeling of I-O target probability

The random forest models and predictions were performed using the “randomForest (v4.6-3)” and “caret (v6.0-93)” libraries in R statistical software. The number of trees in each model was fixed at 1000, as any increase in trees above this value did not yield any significant changes in the prediction probability. The tuneRF function in the random forest package was used to determine the optimal number of variables (“mtry”) sampled at each split. The stepfactor was set at 0.01 and the “improve” parameter at 0.01 value. A positive training set of 45 preclinical and clinical I-O therapeutic targets were identified from the literature (Marshall and Djamgoz 2018). To balance the training set for the model, size-matched negative training sets were generated by random sampling without replacement of genes and excluding the positive training set. Genes with more than 6 missing values were omitted from the predictions. A total of 10,000 random forest models using each of the random negative sets were trained and predictions were made based on each model, followed by averaging the predictions over 10,000 models to assign the IO target probability score to each gene. Model performance was assessed by leave-one-out cross validation using the positive set and a negative set of 185 genes identified by taking the union of non-essential genes from Hart et al (Hart et al. 2014) and those not expressed in lymph node, bone marrow, thymus, whole blood, lymphocytes, and mammary tissues, based on the Human Protein Atlas and GTEx. Predictions were performed using the positive set and a size-matched random sample of the negative controls. The AUC values were averaged over 100 models and the ROC plot was generated using the pROC package from R.

The following features were included in the random forest model:

- *Mendelian genetics* – The Online Mendelian Inheritance in Man^®^ (OMIM^®^) database (https://omim.org/) was mined using Natural Language Processing (NLP) and ontology-based Text and Data Mining (TDM) to extract immunology and cancer phenotype relevant concepts. These data were then converted to a numerical matrix that was compatible for random forest modeling. The full OMIM was obtained via FTP (Morbid Map, Gene Map) and API (full records) respectively (retrieval date: 2022-09-22) and pre-processed with Python for TDM indexing. The information was mined from specific sections (depending on the concept) in the disease and gene monographs, as well as clinical synopses using IQVIA/Linguamatics I2E platform ([CSL STYLE ERROR: reference with no printed form.]). We further utilized internally built knowledge extraction pipelines and disambiguation routines (Alsheikh et al. 2022). TDM queries were focused on paragraphs with clinical / human focus. Linguistic context (sentiment analysis) was used to filter out hits related to normal phenotype (“normal leukocyte count”) or absence of abnormality (“no autoantibodies detected”). TDM identified the most specific concept in the text as hit. In post-processing, ontological hierarchies are considered, and matches propagated to higher level concepts to balance the curator-dependent level of granularity found in OMIM text. For TDM, a list of concepts for normal and pathological immunity were mapped, including those related to immune cells, immunoglobulin types, autoantibodies, anatomical parts of the immune system like spleen, autoimmunity, immunodeficiency, recurrent infection, hypersensitivity, atopy, inflammatory reaction, immune system diseases, leukocyte disorders, and lymphatic diseases (see column headers in **Table S4** for full list of concepts). Likewise, TDM was used to generate a list of concepts for cancer relevant processes (e.g., proliferation, apoptosis) for solid and hematological tumors (**Table S4**). The concepts were mapped to public standard life science ontologies ([CSL STYLE ERROR: reference with no printed form.], [CSL STYLE ERROR: reference with no printed form.], [CSL STYLE ERROR: reference with no printed form.]), and internally enriched with additional synonyms and custom-built vocabularies for concepts that could not be mapped to external standards. Disease concept associations resulting from mining the disease monographs and clinical synopses were turned into indirect Gene – I-O concept associations using enriched OMIM Gene – Disease (G-D) relationships. These relationships use Morbid Map data (6862 G-D relationships) as a basis and are enriched with TDM (5895 additional G-D associations) by mining ‘Molecular Basis’ in clinical synopses and linguistic relationship mining in relevant disease paragraphs. TDM enrichment is crucial as, for example, the association between LSP1 and Neutrophil Actin Dysfunction is only described in the text but not covered in Morbid Map and no indirect LSP1 and I-O associations could have been derived. Finally, all individual direct or indirect gene I-O concept associations were summarized into the binary per gene I-O association matrix (see **Table S4)**. The final OMIM input feature to the random forest model is the sum of concept hits for each gene.
- *Genetic perturbation data* – Published genomewide CRISPR knockout (KO) screens various BrCa cells (Boehm et al. 2021) and immune cell phenotypes ([CSL STYLE ERROR: reference with no printed form.]; Shifrut et al. 2018; Lawson et al. 2020; [CSL STYLE ERROR: reference with no printed form.]; Covarrubias et al. 2020) were curated from the literature. For mouse screens, the mouse genes with human orthologues were converted to the human gene symbols using Ensembl Biomart (Smedley et al. 2015). All screening data are provided in the supplement in the original format and standardized format prior to random forest analysis, as follows: z score = (value – mean) / (standard deviation). Genomewide gene essentiality scores across 40 human BrCa lines (n=10 ER^+^, n=30 ER^-^) in the DepMap (downloaded 4Q19). Essentiality scores were calculated by the DepMap consortium using CERES (Meyers et al. 2017) and averaged across ER^+^ and ER^-^ negative BrCa subtypes (**Table S5-S7**). Genomewide CRISPRi/a screens of the modifiers of IFNg, TNF, or IL2 production in TCR-stimulated human T-cells ([CSL STYLE ERROR: reference with no printed form.]). CRISPR data were analyzed by the authors using MAGeCK version 0.5.9.2. Table column headers are defined as: (A) Gene = Gene name; (B) Screen_Version = Primary CRISPRa/CRISPRi in CD4/CD8 T cells or supplementary screens in CD4 T cells; (C) CRISPRa_or_i = CRISPRa or CRISPRi screen; (D) CD4_or_CD8 = Screen in CD4 or CD8 T cells; (E) Cytokine = Cytokine screened for; (F) LFC = sgRNA median of Log2FoldChange of High/Low sorting bin counts; (G) zscore = Z-Score of LFC values (CRISPRa screens) or −1*LFC (CRISPRi screens), such that positive regulators will have positive Z-Scores and vice versa; (H) FDR = Gene FDR (MAGeCK RRA test); (I) Hit = Hit criteria met (FDR < 0.05 & absolute LFC > 0.5); (J) Hit_Type = Positive or Negative regulator (**Table S8-S9**). Genomewide CRISPR KO screen of CD8+ T-cell expansion in response to TCR stimulation (Shifrut et al. 2018). Significant guide enrichment and depletion was analyzed using MAGeCK and alpha-robust rank aggregation (RRA) to obtain gene-level scores, p-value, FDR, and log fold change (lfc) (**Table S10-S11**). DrugZ ouput summary from genomewide CRISPR KO screens performed in mouse 4T1 and EMT6 breast carcinoma cell lines propagated in the presence or absence of CTLs (Lawson et al. 2020). DrugZ v1 was used to quantitate the fitness effect of gene perturbation under CTL selection. Positive NormZ score indicates gene perturbation results in resistance, whereas negative NormZ score indicates enhanced sensitivity. (**Table S12-S13**). Genomewide CRISPR KO screen of LPS induced NFKB activity in an immortalized bone-marrow-derived macrophage (iBMDM)-Cas9 with an NF-KB-GFP transgenic reporter (Covarrubias et al. 2020). Fold enrichment in LPS-stimulated iBMDM cells was calculated with the lowest Mann-Whitney U test p value (using a p value cut-off of < 0.01) (**Table S14-S15**). Genomewide CRISPR screen in mouse BMDC to identify genes that control the induction of tumor necrosis factor (Tnf) by bacterial lipopolysaccharide (LPS) (Parnas et al. 2015). Differential expression (DE) analysis of the genome-wide screen in “Tnflo” relative to “Tnfhi” was performed using DESeq2. Rank (column E) based on the p-value (include positive and negative regulators). Standard DESeq output (columns F–K), including mean expression of all guides targeting each gene (column F) and the fold change (column G) between “Tnflo” and “Tnfhi.” Positive values indicate enrichment in the “Tnflo” library and therefore positive regulators. Negative values indicate depletion in the “Tnflo” library and therefore negative regulators (**Table S16-S17**). Genomewide CRISPR screen in TH2-differentiated T-cells that were isolated from transgenic mice expressing Cas9 and fluorescent reporters for Il4, Il13, Xbp1, and Gata3. LogFC are reported at gene level for guides enriched in high vs low populations at 72h post differentiation to TH2 phenotypes (Henriksson et al. 2019). Data were analyzed by BaIOPSE and MAGeCK, and reported at gene level as logFC, p-value, and ranked score (**Table S18-19**).
- *MGI Mouse Knockout Phenotypes* – Gene annotations for immunological and BrCa phenotypes from genetically engineered mouse models (GEMMs) were downloaded from the MGI database (Bult et al. 2019) (**Table S20-S21**).
- *COSMIC Cancer Gene Census* – Cancer driver gene annotations for BrCa were queried from Cancer Gene Census list in the COSMIC database (Sondka et al. 2018) (**Table S22**).
- *Leukocyte eQTL* – Genes annotated were annotated for significant eQTL (FDR < 0.05) across 28 immune cell types from 416 donors in the ImmuNexUT database (Ota et al. 2021). Effect sizes for significant associations were used as inputs for the random forest model (**Table S23**).

### LSP1 transcription expression in human breast cancer biopsies by scRNAseq analysis

The BioTuring BBrowser3 was used to visualize cell type clusters and LSP1 transcript expression from 14 treatment-naïve breast cancer patient RNAseq dataset with a total of 44,024 cells profiled (Qian et al. 2020). Differential expression analysis of LSP1 was performed using the BioTuring package “Venice”, as described previously (Le et al. 2020), comparing LSP1 expression in T/NK population (14,395 cells) to cancer population (16,235 cells) as defined by the BioTuring software cell-type specific markers.

### LSP1 protein expression in human breast cancer tissue biopsy microarrays

A validated multiplexed immunohistochemistry protocol was used to detect LSP protein in tumor-associated leukocytes in 1,109 de-identified cases of invasive BrCa diagnosed at the Jefferson University Hospital from 1988-2005 and made available under institutional IRB approved protocols in tissue microarray format (0.6 mm core diameter) as previously described (Andersson et al. 2021; Peck et al. 2016). In addition to LSP1, multiplex fluorescence staining also included CD45 to detect leukocytes and pan-cytokeratin to detect cancer cells. Immunohistochemistry was performed on an autostainer (Omnis; Agilent/Dako). After antigen retrieval at pH 6, TMA sections were incubated with polyclonal rabbit anti-LSP1 antibody (Sigma HPA019693; 1:8000 dilution), followed by HRP-conjugated anti-rabbit secondary antibody polymer (EnVision+; Dako-Cat#K4003), and visualized using Cy5-tyramide as substrate. The antigen retrieval step was repeated followed by second round staining for pan-CK using anti-pan-cytokeratin antibody (AE1/AE3 mouse monoclonal, Dako-Cat# M351501-2) followed by HRP-conjugated anti-mouse secondary antibody polymer (EnVision+; Dako-Cat# K400311-2) and visualized by Cy7-tyramide as substrate. CD45 was stained using biotin-conjugated anti-CD45 (Biolegend; cat#304004; 1:50 dilution) and visualized by incubation with Alexa555-labeled streptavidin. DAPI counterstain was used to visualize cell nuclei. Stained slides were digitized at 20x magnification on a slide scanner (3DHistech Pannoramic Flash II) capturing fluorescent images captured in four channels (DAPI, Alexa555, Cy5, ad Cy7. Digitized images were analyzed by Tissue Studio (Definiens) and mean cytoplasmic expression signal for LSP1 immunoreactivity was computed for CD45-positive cells in each tumor core. All immunohistochemistry, slide scanning, and quantitative analyses of digitized images were performed by investigators that were blinded to outcome.

### Analysis of patient outcome associated with LSP1 protein expression

Clinical outcome data (progression-free survival; PFS, median follow-up 8.8 years) were available for the 1,109 ER+ BrCa patients. A data-driven optimal cutpoint for dichotomization (High vs. Low) of the LSP1 expression in CD45-positive cells was derived using X-tile (Camp et al. 2004) and the prognostic value of the resulting dichotomized biomarker was evaluated using the Log-rank test and Kaplan-Meier curves (IBM SPSS).

### Animal models

All animal experiments were performed at the Medical College of Wisconsin (MCW) and all protocols were approved by the Institutional Animal Care and Use Committee of MCW. PWD/PhJ, C57BL/6J, and B6.FVB-Tg(MMTV-PyVT)634Mul/LellJ mice were purchased from Jackson Laboratory and maintained at MCW. The F1 and N1 strains for phenotyping were generated as follows: PWD/PhJ x C57BL/6J (PWD/C57), C57BL/6J x C57BL/6J (C57/C57), PWD/PhJ x B6.FVB-Tg(MMTV-PyVT)634Mul/LellJ (PWD/C57-PyVT), and C57BL/6J x B6.FVB-Tg(MMTV-PyVT)634Mul/LellJ (C57/C57-PyVT).

### Spontaneous mammary tumor model and linkage analysis

The F_1_ generations of the PWD/C57-PyVT and C57/C57-PyVT crosses were genotyped for the PyVT transgene using an established protocol from the Jackson Laboratory. Beginning at 6 weeks-of-age, the transgene-positive PWD/C57-PyVT and C57/C57-PyVT female mice were palpated weekly for mammary tumors, and the initial date of detection and anatomical location were recorded. Prior to tissue collection, tumor-positive mice were then aged for 40 days postdiagnosis to permit development of metastases. To localize the mammary tumor modifiers in the PWD/PhJ and C57BL/6J genomic backgrounds, a [C57BL/6J x (PWD/PhJ x B6.FVB-Tg(MMTV-PyVT)634Mul/LellJ)]N2 backcross (n = 104) was performed. Genome-wide genotyping was performed using 3,753 informative markers from the MUGA array (Morgan et al. 2015). A single-locus scan was performed and LOD scores were calculated at 0.5 cM interval across the genome, using the imputation method implemented in R/qtl (Sen and Churchill 2001) and significance was determined on the basis of 1,000 permutations of the data (Churchill and Doerge 1994). A LOD score exceeding the 0.1 genome-wide adjusted threshold was considered significant (Lander and Kruglyak 1995). The Bayes credible interval function in R/qtl (bayesint) was used to approximate the 95% confidence intervals for the QTL peak location for both the additive and the interactive models, as described in (Solberg et al. 2004). Mouse QTL boundaries were converted from cM to Mb using the Mouse Map Converter tool (http://cgd.jax.org/mousemapconverter/), and genomic features and variant effect predictions (VEP) were annotated using the “Genes and Markers Query” and “Search Mouse SNPs” functions of the Mouse Genome Informatics (MGI) database (http://www.informatics.jax.org/).

### Mammary tumor transplant model

EO771.LMB cells were orthotopically implanted into PWD/PhJ x C57BL/6J and C57BL/6J x C57BL/6J F1 animals and measured as described previously into (Lifsted et al. 1998), with slight modifications. Briefly, EO771.LMB cells (5 × 10^5^) in 50% Matrigel were implanted into the mammary fat pad (MFP) of age-matched female PWD/PhJ x C57BL/6J F_1_ (n = 10) and C57BL/6J x C57BL/6J N1 (n = 22) mice. Tumor volumes were measured weekly by calipers, as described previously (Whitehurst et al. 2007).

### RNAseq library preparation and sequencing

Total RNA was collected from EO771.LMB tumors that were xenografted into B6 and PWD.B6 mice (n = 4 per group) isolated using TRIzol Reagent (ThermoFisher), poly-A purified, transcribed, and chemically fragmented using Illumina’s TruSeq RNA library kit using the manufacturer’s protocol. Individual libraries were prepared for each sample, indexed for multiplexing, and then sequenced on an Illumina HiSeq 2500 (Illumina, Inc., San Diego, CA). Genome sequence and GTF files were obtained from Ensembl. The RSEM (RNA-Seq by Expectation-maximization) program function “rsem-prepare-reference” (v1.3.0) was used to extract the transcript sequences from the mouse genome (build GRCm38) (Li and Dewey 2011) and to generate Bowtie2 indices (Bowtie2 v2.2.8) (Langmead and Salzberg 2012) followed by read alignment using the “rsem-calculate-expression” function. Differential expression analysis was performed using the Bioconductor package DESeq2 version 1.16.1 (Love et al. 2014) to compute log2 fold-changes and FDR-adjusted *P* values. Molecular and functional pathway enrichment was measured using the Ingenuity Pathway Analysis (IPA) tool (QIAGEN, Redwood City, CA). Gene set enrichment analysis (GSEA) was performed as described previously (Subramanian et al. 2005; Mootha et al. 2003). Enrichment of tumor infiltrate signatures was estimated using single-sample GSEA (ssGSEA) using a compiled list of immune 23 signatures (Bindea et al. 2013; Hendrickx et al. 2017; Martinez et al. 2015; Monaco et al. 2019), as implemented by the R package “GSVA” (v 1.34.0) and following ortholog mapping between human and mouse genes using the “biomaRT” R package.

## DATA ACCESS

The supporting data are available as Supplemental Tables.

## COMPETING INTEREST STATEMENT

B.R.G., E.K., S.W., J.R., Z.D., and M.J.F. are employees of AbbVie. All animal studies and histological analysis of human breast cancer specimens were conducted at the Medical College of Wisconsin (MCW), at which time M.J.F was a full-time faculty member of MCW. S.W.T., A.R.P., K.W., A.L., and H.R. are employees of MCW and have no financial relationship with AbbVie to disclose. The design, study conduct, and financial support for all other research were provided by AbbVie. AbbVie participated in the interpretation of data, review, and approval of the publication.

## ACKNOWLEDGEMENTS

We thank AbbVie employees Cyril Ramathal, Lynn Miller, and Meagan Fricano for the helpful discussions and critical feedback. We thank the AbbVie Research and Development Convergence Hub (ARCH) for curation of genomics datasets. We thank MCW employee Linna Ge for expert immunohistochemistry, no funding to disclose. This work was supported, in part, by funding from grants from the Wisconsin Breast Cancer Showhouse (M.J.F. and H.R.) and the MCWCC, the Advancing a Healthier Wisconsin Endowment [no. 5520367 (H.R.) and no. 5520423 (M.J.F. and H.R.)], the Dr. Nancy Laning Sobczak Fund for Breast Cancer (H.R. and M.J.F.). Support was also received from the NCI [R01CA193343 (M.J.F.), R01CA188575 (H.R.)], the Mary Kay Foundation [grant no. 024-16 (M.J.F.)], and the METAvivor Foundation (M.J.F. and H.R.). We acknowledge and thank the participants and investigators of the UK Biobank, this research was carried out using the UK Biobank resource under application number 26041. UK Biobank was established and funded by the Wellcome Trust medical charity, Medical Research Council, Department of Health, Scottish Government and the Northwest Regional Development Agency. UKBB has also received funding from the Welsh Government, British Heart Foundation, Cancer Research UK and Diabetes UK. UK Biobank is supported by the National Health Service (NHS). We want to acknowledge the participants and investigators of the FinnGen study. The FinnGen project is funded by two grants from Business Finland (HUS 4685/31/2016 and UH 4386/31/2016) and the following industry partners: AbbVie Inc., AstraZeneca UK Ltd, Biogen MA Inc., Bristol Myers Squibb (and Celgene Corporation & Celgene International II Sàrl), Genentech Inc., Merck Sharp & Dohme LCC, Pfizer Inc., GlaxoSmithKline Intellectual Property Development Ltd., Sanofi US Services Inc., Maze Therapeutics Inc., Janssen Biotech Inc, Novartis AG, and Boehringer Ingelheim International GmbH. Following biobanks are acknowledged for delivering biobank samples to FinnGen: Auria Biobank, THL Biobank, Helsinki Biobank, Biobank Borealis of Northern Finland, Finnish Clinical Biobank Tampere, Biobank of Eastern Finland, Central Finland Biobank, Finnish Red Cross Blood Service Biobank, Terveystalo Biobank and Arctic Biobank.

